# Migration rather than proliferation transcriptomic signatures are strongly associated with breast cancer patient survival

**DOI:** 10.1101/379123

**Authors:** Nishanth Ulhas Nair, Avinash Das, Vasiliki-Maria Rogkoti, Michiel Fokkelman, Richard Marcotte, Chiaro G. de Jong, Joo Sang Lee, Isaac Meilijson, Sridhar Hannenhalli, Benjamin G. Neel, Bob van de Water, Sylvia E. Le Dévédec, Eytan Ruppin

## Abstract

The efficacy of prospective cancer treatments is routinely estimated by *in vitro* cell-line proliferation screens. However, it is unclear whether tumor aggressiveness and patient survival are influenced more by the proliferative or the migratory properties of cancer cells. To address this question, we experimentally measured proliferation and migration phenotypes across more than 40 breast cancer cell-lines. Based on the latter, we built and validated individual predictors of breast cancer proliferation and migration levels from the cells’ transcriptomics. We then apply these predictors to estimate the proliferation and migration levels of more than 1000 TCGA breast cancer tumors. Reassuringly, both estimates increase with tumor’s aggressiveness, as qualified by its stage, grade, and subtype. However, predicted tumor migration levels are significantly more strongly associated with patient survival than the proliferation levels. We confirm these finding by conducting siRNA knock-down experiments on the highly migratory MDA-MB-231 cell lines and deriving gene knock-down based proliferation and migration signatures. We show that cytoskeletal drugs might be more beneficial in patients with high predicted migration levels. Taken together, these results testify to the importance of migration levels in determining patient survival.

## Introduction

Drug development risk is a major contributing factor for spiraling drug prices^1^. Only 1 out of 5000 drugs from pre-clinical studies enter the market after successful clinical testing^2^. Cancer drugs show the highest proportion of failures on the road to clinics^3^. Currently, the prevailing experimental method to initially estimate the pre-clinical efficacy of cancer drug candidates is by measuring their effects on *in vitro* proliferation rates^3–10^. However, even after filtering these findings in animal models, only a fraction of emerging candidates has successfully translated into human trails^11–13^. Many factors contribute to the failure of drugs that are effective in pre-clinical systems. For starters, *in vitro* and *in vivo* systems are obviously only approximate models of patients that do not capture many aspects of human biology. However, another naturally arising possibility is that other cellular phenotypes, such as migration or invasion, may be better indices of tumor response in patients than cellular proliferation. Addressing this question, we aimed here to quantify the relative weight of proliferation verses migration in determining cancer aggressiveness and patient survival.

Ideally, one would have liked to directly measure proliferation and migration levels directly in tumors *in vivo* to study their association with patient survival and treatment response. However, regrettably, such measurements are yet infeasible. We therefore set out to build and validate predictors of proliferation and migration levels in breast cancer cell-lines based on gene expression information. Having such predictors in hand, we then apply them to TCGA breast cancer tumors to predict the levels of these phenotypes in these tumors from their gene expression. We tested and verified that the predicted levels of these phenotypes in the tumors are indeed strongly associated with patient survival in the direction expected, and that they are associated with cancer aggressiveness as expected. We find that migration is more strongly associated with breast cancer aggressiveness and more importantly, patient survival, than proliferation.

## Results

### Overview

We built predictors of cell proliferation and migration as follows: First, we experimentally measured migration and proliferation values in 43 and 46 breast cancer cell lines respectively (Table S1a, Methods). Second, we constructed gene-expression based predictors of migration and proliferation, termed *CellToPhenotype* predictors, using least absolute shrinkage and selection operator (LASSO) based regression^14^. The predictors were tested on the cell-line data using a standard cross-validation procedure (Methods, Table S1b). Third, we used the predictors built to estimate the migration and proliferation levels of 1043 breast cancer patients in TCGA data (Table S1c). Finally, we explored their importance in predicting tumor stage, grade, subtypes, and patients’ survival (see Figure 1 for an overview).

**Figure 1:**
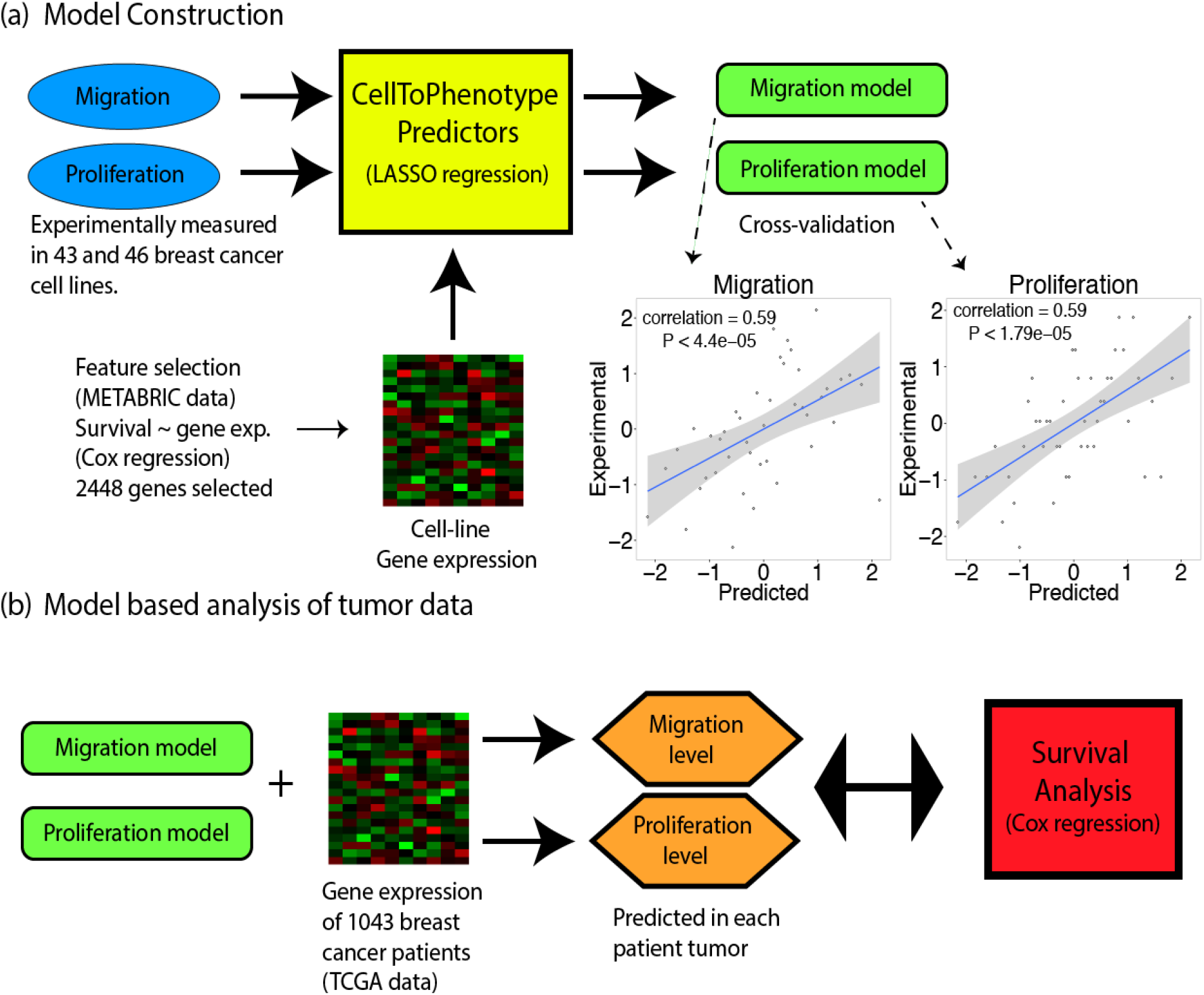
Overview of the method. **(a)** CellToPhenotype predictors of migration and proliferation from gene expression are constructed from experimentally determined migration and proliferation measurements across 43 and 46 breast cancer cell lines respectively. The predictors are built using cross-validation, and the correlations obtained between predicted levels and actual experimentally measured values are depicted as scatter plots. **(b)** The CellToPhenotype predictors are used to analyze the gene expression values of breast cancer patients to predict migration and proliferation levels of 1043 TCGA breast cancer tumors. Subsequently, the association of tumors predicted migration and proliferation levels with different tumor phenotypes and patients’ survival is examined.

### Experimental measurements of migration and proliferation

Doubling times for 46 breast cancer cell lines were estimated by plating a known number of cells and measuring the total number of cells once the culture reached an estimated 80% confluency. Proliferation rates were then calculated using the doubling time measurements (Methods). Cell line migration was estimated in 43 breast cancer cell lines using a live cell migration assay. The mean speed of cell migration was then quantified as the final migration estimate (Methods).

### *CellToPhenotype* predictor construction and validation

We measured migration and proliferation values in a collection of breast cancer cell lines available to us, for which we also had the transcriptomics data of each cell-line (Table S1a, Methods). Based on this data, we constructed predictors of migration and proliferation, termed *CellToPhenotype* predictors, which given the expression of a given cell-line, predict its migration and proliferation levels. These predictors were constucted using least absolute shrinkage and selection operator (LASSO) based regression^14^, considering as features the genes whose expression is significantly associated with survival in the METABRIC breast cancer collection (Methods). The CellToPhenotype predictors accurately estimate cell-line migration (Spearman ρ=0.59, P<4.39e-5) and proliferation values (Spearman ρ=0.59, P<1.79e-5) using a standard cross validation procedure (Figure 1a). The genes selected as the features used by the CellToPhenotype proliferation and migration predictors are shown in Table S3(a-d). These features sets are enriched for RAC1 signaling pathway (RAC1 is associated with cell motility^15^), immune response and cell apoptosis (Table S3(e,f)) in the migration predictor. The gene features of the proliferation predictor are enriched in cell differentiation, promoter transcriptional regulation and tissue development (Table S3(g,h)). A KEGG pathway analysis of these genes shows an enrichment in cancer related pathways known to be involved in migration and proliferation. These include HIF-1 signaling and ECM receptor interaction for migration and ErbB signaling and transcriptional misregulation for proliferation (Table S3i, Methods).

Next, we compared CellToPhenotype predictions with expression of three reported gene markers of migration and proliferation. First, the expression of Ki-67, a known marker of cell proliferation and patient survival^16–18^. Its expression is correlated with the experimental cell line proliferation measurements is significant but weaker (Spearman ρ = 0.32, P<0.03) than that obtained by the CellToPhenotype predictor. Second, as a control, MIB-1 expression, a marker of tumor cell proliferation and determinant of patient survival in prostate cancer^19^, shows no correlation with experimental measured proliferation in breast cancer cell lines (Spearman ρ = 0.05, P<0.76). Finally, TPX2 expression, a known correlate of migration in breast cancer^20^, is significantly correlated (Spearman ρ = 0.4, P< 0.008) with the experimental measurements of migration in breast cancer cell lines. Again, this correlation is smaller compared to correlation via the CellToPhenotype migration predictor. Overall, these results show that the latter provide a better estimate of *in vitro* proliferation and migration than known marker genes.

### Predicted migration and proliferation levels are significantly higher in tumor samples than in normal samples

We then applied the CellToPhenotype proliferation and migration predictors to analyze breast cancer TCGA tumor data. Given an input tumor sample, each predictor (migration or proliferation) receives as input the levels of expression of its feature genes in that sample, and outputs the predicted migration or proliferation levels. We first tested if the CellToPhenotype predicted proliferation and migration levels are higher in the TCGA breast tumors than matched adjacent tissues (analyzing 110 TCGA breast cancer patients for which such matched data exists). Reassuringly, the predicted migration and proliferation levels are significantly higher in the tumors than in the matched noncancerous tissues (paired Wilcoxon rank-sum test^21^, P<5.5e-20 and P<4.4e-20 respectively, Figure 2a). Random linear combinations of survival-significant genes used as control predictors do not show any such differences either for migration or proliferation (paired Wilcoxon rank-sum test, P<0.6 and P<0.47 respectively, Supplementary note).

**Figure 2:**
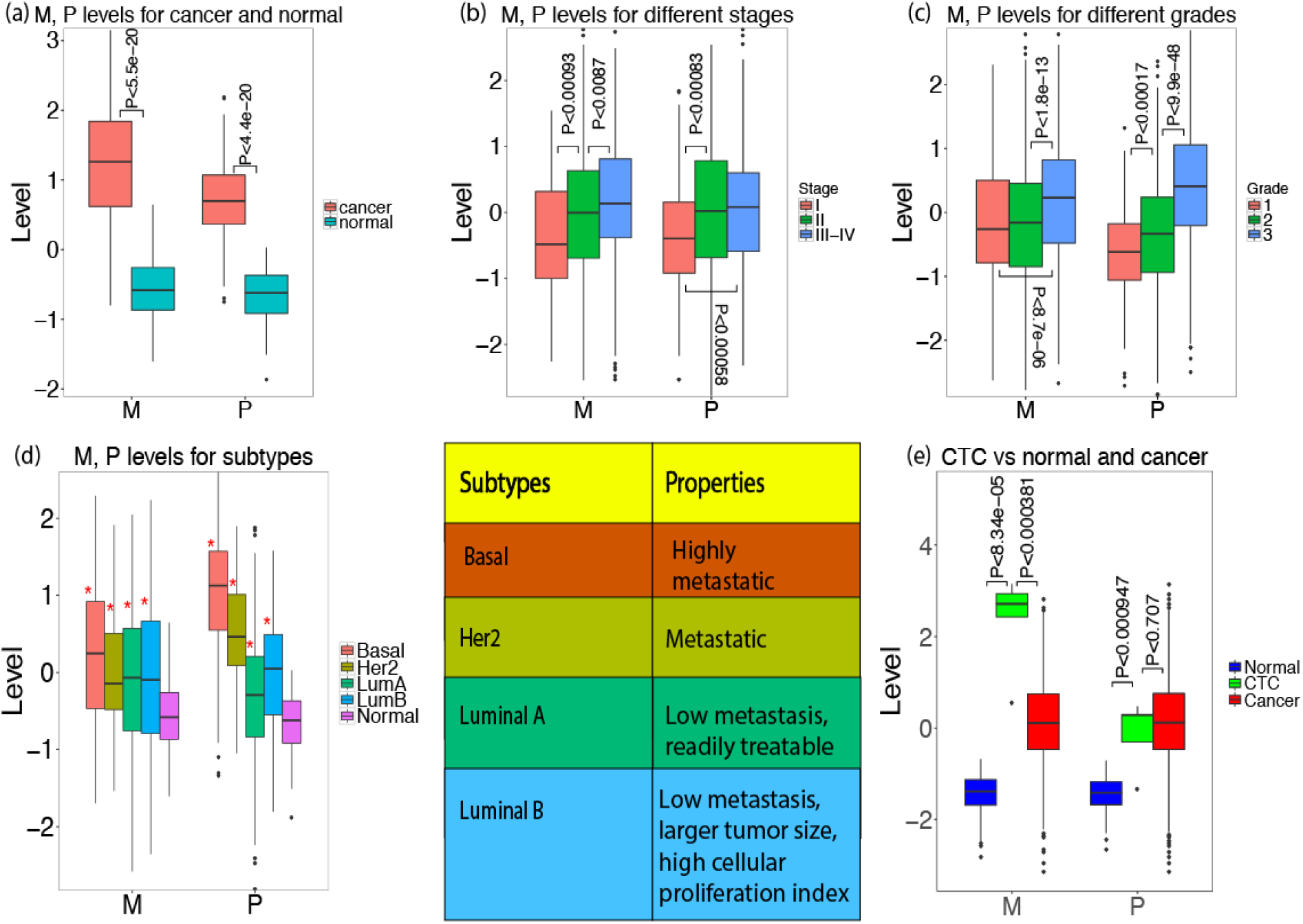
**(a)** Predicted migration (M) and proliferation (P) levels of breast cancer tumors and their association with various clinical phenotypes. (a) M, P levels of 110 breast cancer patients for their tumor (cancer) and matched non-cancerous breast samples (normal). **(b)** Predicted M, P levels for 937 breast cancer TCGA tumors for which cancer stage information is available. **(c)** Predicted M, P levels for 1706 METABRIC breast cancer patients for which cancer grade information is available. **(d)** Predicted M, P levels for 497 breast TCGA tumors dataset having subtype information: Basal or Triple-Negative (91 patients), Her2 (55 patients), Luminal A (LumA, 224 patients), Luminal B (LumB, 127 patients), and noncancerous samples (110 patients). The properties of these subtypes are shown in a table. Significant differences (when comparing tumors to non-cancerous samples) are marked via ‘*’. **(e)** Predicted M, P levels of 5 samples of circulating tumor cells (CTC) from GSE45965 data, compared with the 110 noncancerous samples and 1043 breast cancer TCGA samples.

### Advanced stages of breast cancer have higher predicted migration and proliferation levels than early stages

We next tested if CellToPhenotype predicted migration and proliferation levels would be higher in advanced stage patients, as expected. 937 breast cancer patients have cancer stage information available in TCGA (Table S2a). Indeed, the predicted migration levels increased significantly from stage I to stage II (Wilcoxon rank sum test, P<9.3e-4); and from stage II to stage III-IV (P<8.7e-3). Predicted proliferation levels also increase from stage I to stage II (P<8.3e-4) and from stage I to stage III-IV (P<5.8e-4) (Figure 2b, Methods). As a control, random linear combinations of survival-significant genes do not exhibit any significant increase in their levels with higher tumor stages (Supplementary note).

### Predicted migration and proliferation levels increase with cancer grade

We next tested if the predicted migration and proliferation increase with cancer grade. Tumor grade information was available for 1706 breast cancer patients in METABRIC (Grade information is absent in TCGA breast cancer patients; Methods, Table S2a). Predicted migration levels were indeed significantly higher in grade 3 patients when compared to grade 2 (Wilcoxon rank-sum P<1.8e-13) and grade 1 patients (P<8.7e-06). Similarly, the predicted proliferation levels increase significantly from grade 1 to grade 2 (P<1.7e-4), and from grade 2 to grade 3 patients (P<9.9e-48) (Figure 2c). Random linear combinations of survival-significant genes do not exhibit any such association with tumor grade (Supplementary note).

### Predicted migration and proliferation levels match known attributes of breast cancer subtypes

Different breast cancer subtypes have been associated with different migration and proliferation phenotypes. We therefore asked whether the predicted levels using CellToPhenotype predictors recapitulate these attributes. We find that all four subtypes of breast cancer (Table S2a; basal or triple negative breast cancer (TNBC), Her2, Luminal A and B) have significantly higher migration and proliferation levels than that of non-cancerous samples (n=110) (Figure 2d, Methods). Consistent with the observation that TNBC tumors are highly metastatic^22^, we find that TNBC patients (n=91) exhibit the highest predicted migration and proliferation levels amongst subtypes. Luminal A patients (n=224) exhibit the lowest predicted proliferation and migration levels amongst subtypes, consistent with the observation that they have low metastasis levels and respond relatively well to treatment. Luminal B (n=127) patients have higher predicted proliferation levels than Luminal A patients, consistent with the observation that Luminal B has larger tumor size and higher cellular proliferation index^22^, and higher rates of lymph node involvement than Luminal A^23^ (Figure 2d).

### Circulating tumor cells have high migration levels

We applied the CellToPhenotype predictors to predict the migration and proliferation levels in 5 samples of circulating breast tumor cells (CTCs) in GSE45965 data^24^. We find that CTC samples have significantly higher migration levels than both the TCGA cancer samples (P<3.81e-4) and the healthy adjacent samples (Wilcoxon rank sum, P<8.34e-5). The CTC samples have significantly higher proliferation levels than the non-cancerous samples (P<9.47e-4), but not significantly higher than the cancer samples (P<0.71, Figure 2e).

### Predicted migration levels are more strongly associated with patient survival than predicted proliferation levels

We studied the association of predicted migration and proliferation levels with patients’ survival in the TCGA breast cancer dataset (1043 patients). For this analysis, we first built new predictors of migration and proliferation by analyzing the 40 breast cancer cell lines for which we have both migration and proliferation measurements to enable a head-to-head comparison of the effects of migration and proliferation on survival, when built from exactly the same cell-lines (the construction itself followed the same predictor generation procedure described above). Given these CellToPhenotype predictors we estimated the migration and proliferation levels of the 1043 TCGA breast tumors from their expression data, as before. We then employed a Cox regression to examine the association between these predicted values and the patients’ survival, controlling for various confounders including age, race, and genomic instability (Methods). We find a stronger association of predicted migration levels with patient survival (risk factor = 0.45, P<2.06e-5) than the association between proliferation and survival (risk factor = 0.36, P<8.25e-4) (Figure 3a, Methods). In both cases, the higher the predicted migration/proliferation levels are, the lower is patients’ survival. A similar trend is revealed using a Kaplan Meier (KM) analysis (Kaplan and Meier, 1958) comparing tumors with high vs low predicted migration and proliferation levels (Figure 3b). A multivariate Cox-regression (Methods, Figure 3c) again shows that migration is more strongly associated with patient survival (relative risk factor = 0.35, P<2.82e-3) than proliferation (relative risk factor = 0.20, P<0.102). Random linear combinations of survival-significant genes do not exhibit any significant associations with patient survival (risk factor = 0.067, P<0.24 for migration; risk factor = −0.0021, P<0.24, for proliferation; Supplementary note).

**Figure 3:**
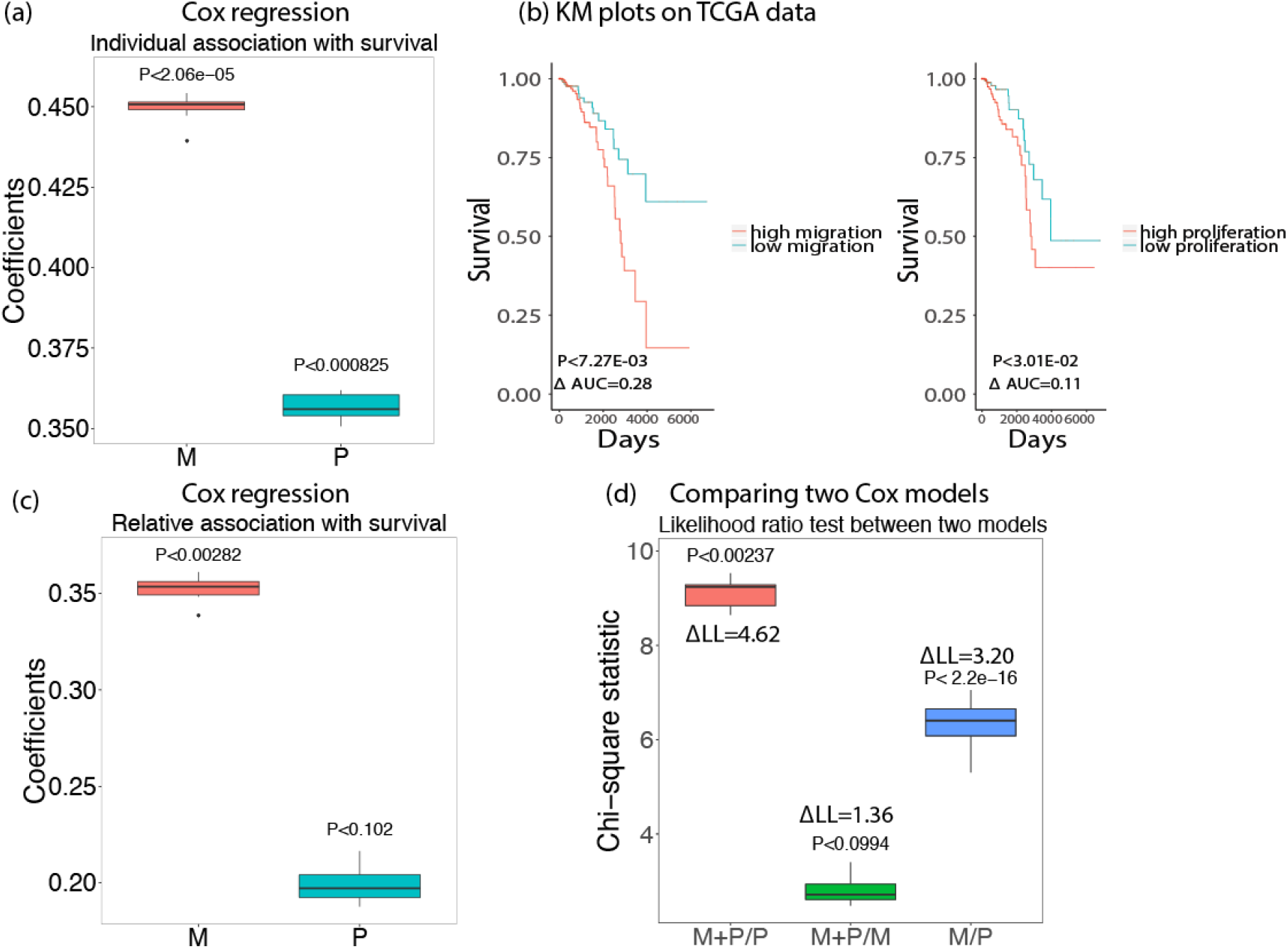
Survival analysis for 1043 breast cancer patients in TCGA data using predicted migration (M) and proliferation (P). Box plots of 10 iterations are shown and median p-value of each coefficient is given above the box plots. A positive coefficient (risk factor) for M (or P) indicates that the higher value of M (or P), the lower the patient survival. **(a)** Coefficients of M, P when used to predict survival individually using Cox regression after controlling for age, race, and genomic instability. **(b)** A Kaplan Meyer (KM) survival analysis of tumors’ predicted migration and proliferation levels. **(c)** Relative coefficients of M, P when used to predict survival when they are controlled by each other (multivariate Cox-regression). **(d)** Likelihood ratio test comparing how significantly different are two Cox regression models with each other (Chi-square test statistics with p-values are provided): (i) Migration and Proliferation vs Proliferation only (M+P/P); (ii) Migration and Proliferation vs Migration only (M+P/M); (iii) Migration only vs Proliferation only (M/P). The difference between log-likelihood (ΔLL) between the two models is also shown.

To estimate significance of difference of predictability, we additionally performed a likelihood ratio test comparing the survival predictive power of combined migration and proliferation compared to only proliferation or only migration. We see that the migration + proliferation model is significantly better than a proliferation only model (Chi-square statistic=9.24, P<2.37e-3) but not significantly better from a migration only model (Chi-square statistic=2.72, P<0.099). Thus, adding migration to a proliferation-only model improve the survival prediction, however adding proliferation does not add significant predictive power to a migration-only model., A migration only model is significantly better than a proliferation only model (Chi-square statistic=6.40, P<2.2e-16) in predicting patient survival (Methods, Figure 3d). Finally, we observed that survival prediction accuracy was considerably reduced if we use a smaller number of cell lines for building the CellToPhenotype predictors (Supplementary note), thus showing the importance of studying a large number of cell lines.

### A siRNA-based analysis further supports that migration is more strongly associated with survival than proliferation

We next turned to study our basic research question by building an additional set of predictors of migration and proliferation levels. These predictors are based on siRNA knockdown (KD) experiments that we have conducted in a highly migratory breast cancer cell line, MDA-MB-231. We knocked down 248 protein kinases whose gene expression levels are significantly negatively correlated with experimentally determined migration values in the 40 breast cancer cell lines that we studied above. We experimentally measured the effect of each knockdown on cell migration using a 2D migration assay (Methods). We termed the genes whose knockdown significantly enhances cellular migration *migration-suppressive genes* (Methods). Similarly, we also carried out siRNA KD experiments on 227 protein kinases whose gene expression levels are significantly positively correlated with experimentally determined migration values in these 40 breast cancer cell lines, and determined the effect of each knockdown on cell migration. Among these, we identified all genes whose knockdown significantly decreased cellular migration in MDA-MB-231 cell line and termed them as *migration-enhancer* genes (Methods). Migration suppressive genes (using the siRNA-based analysis, Table S3j) show enrichment in gene sets involved in metastasis and breast cancer and cell migration (Table S3k). Migration-enhancer genes (Table S3j) show enrichment in actin cytoskeleton, focal adhesion and cell-cell junctions (Table S3(i,m)).

The number of migration-suppressive genes that were downregulated in a given breast cancer cell-line (S-count) was highly correlated with its CellToPhenotype predicted migration levels (Spearman ρ = 0.81, P<3.43e-10) and also with its experimentally measured migration values (Spearman ρ = 0.79, P<1.83e-9). This suggests that the S-count could be considered as an approximation of cellular migration. Similarly, the number of migration-enhancer genes that were upregulated in a given breast cancer cell-line denotes their migration-enhancer scores (E-count). The E-score is also highly correlated with the predicted migration levels (Spearman ρ = 0.827, P<8.62e-11) and the experimentally measured migration values (Spearman ρ = 0.77, P<1.03e-8), suggesting that E-count also provide approximate estimates of cellular migration. The mean of S-count and E-counts, termed the KD-migration-score (Methods), has a slightly higher correlation with the predicted migration levels (Spearman ρ = 0.83, P<4.36e-11, Supplementary Figure S1a) and experimentally measured ones (Spearman ρ = 0.79, P<1.9e-9, Supplementary Figure S1b) across the cell-lines.

Similarly, in an analogous manner, we computed a KD-proliferation-score, using published shRNA/siRNA knockdown data done in MDA-MB-231 cell line^25^ (Methods). The KD-proliferation-scores are highly correlated with both the predicted proliferation levels (Spearman ρ = 0.75, P<3.07e-8, Supplementary Figure S1c) and the experimentally measured proliferation values (Spearman ρ = 0.82, P<2.57e-10, Supplementary Figure S1d). Reassuringly, we find that the cross-correlations between KD-migration-score and experimentally-measured proliferation levels (and vice-versa) are much lower (Supplementary note). Gene Set Enrichment Analysis (GSEA) analysis on proliferation-enhancer/suppressive genes, showed enrichment on interesting gene sets, including, cell proliferation, regulation of developmental processes, genes associated with breast cancer (Table S3(n,o)).

Having these scores in hand, we next computed *KD-migration-scores* and *KD-proliferation-scores* for every TCGA breast cancer tumor. Reassuringly, these scores are significantly correlated with the CellToPhenotype predictions of migration (Spearman ρ = 0.4, P<5.95-42) and proliferation levels of these tumors (Spearman ρ = 0.69, P<2.77-150). We then examined the association of the *KD-migration* and *KD-proliferation scores* of the TCGA breast cancer tumors and patient survival, after controlling for age, race, and genomic instability via Cox regression. The results reinforce the trend observed previously with the CellToPhenotype analysis, as we find a significant association of *KD-migration-scores* with patient survival (risk factor = 0.3, P<2.98e-3) but a lower association between *KD-proliferation-scores* and survival (risk factor = 0.24, P<0.0396) (Figure 4a, Methods). A multivariate cox-regression using both the *KD-migration-scores* and *KD-proliferation-scores* as co-variates while controlling for age, race, and genomic instability shows a similar trend (relative risk factor = 0.29, P<0.037 for migration and relative risk factor = 0.044, P<0.76 for proliferation, Figure 4b, Methods). Thus, ruling out model-based biases of CellToPhenotype predictors, the analysis further corroborates our findings that migration is better predictor patient survival than proliferation. As it is knock-down based, it suggests that the stronger association of migration with survival may have a causal basis.

**Figure 4:**
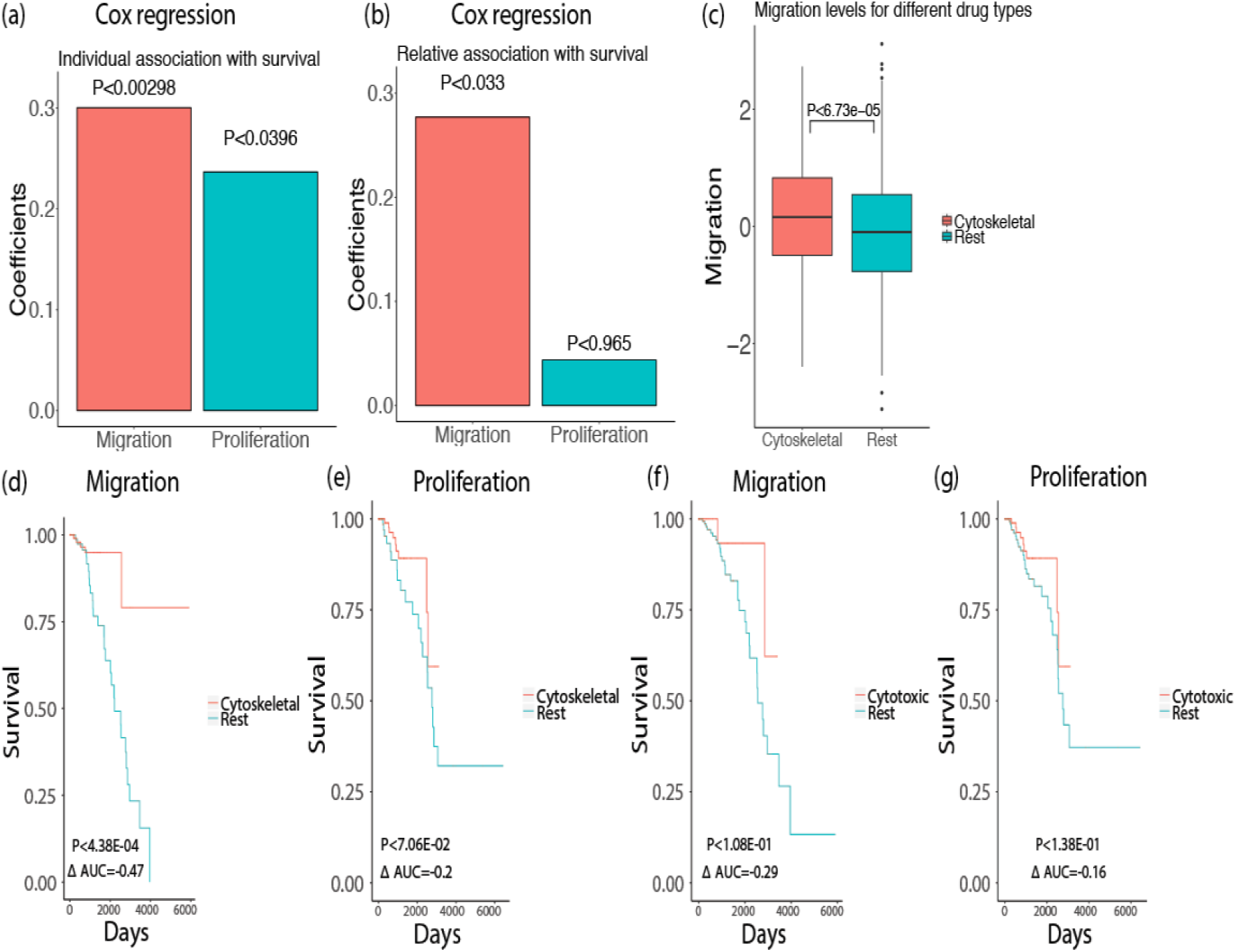
**(a)** Cox regression of KD-migration-scores and KD-proliferation-scores with patients’ survival, after controlling for age, race, and genomic instability for 1043 breast cancer patients in TCGA data. **(b)** Relative association of KD-migration-scores and KD-proliferation-scores with survival. **(c)** Migration levels (estimated from CellToPhenotype predictors) for breast cancer patients who have taken cytoskeletal drugs versus the rest. **(d)** Among the breast cancer patients who have high migration levels (greater than 75 percentile), a KM analysis was done between those who have taken cytoskeletal drugs versus the rest of the patients. **(e)** Similarly, among the patients that have high proliferation levels, a KM analysis was done between those who have taken cytoskeletal drugs versus the rest of the patients. **(f)** A KM analysis of patients who have high migration levels and who have taken only cytotoxic drugs versus the rest. **(g)** A KM analysis of patients who have high proliferation levels and have taken only cytotoxic drugs versus the rest.

### CellToPhenotype estimates of migration and proliferation levels are associated with patient drug response

We next asked if predicted migration or proliferation levels determine patient response to cytoskeletal vs cytotoxic drugs. Although migration and proliferation occur via partially overlapping cellular processes, as a first approximation cytotoxic drugs mainly target proliferation by inhibiting nucleotide synthesis or inducing DNA break while cytoskeletal drugs are known to affect cell migration by targeting microtubules (though they also target proliferation)^26,27^. Accordingly, our working hypothesis has been that tumors with high migratory estimates may respond better to cytoskeletal drugs and conversely, tumors with high proliferation estimates will respond better to cytotoxic drugs.

To test this hypothesis, we analyzed data of TCGA breast cancer patients, out of which 389 patients were given at least one of the 7 cytoskeletal drugs and 331 patients were given at least one of 31 cytotoxic drugs (Table S2b). In the patients treated with cytoskeletal drugs, migration levels (estimated from CellToPhenotype predictors) are significantly higher than the levels in the rest of TCGA breast cancer patients (P<6.73e-5, Figure 4c), and higher than the levels of the patients treated only with cytotoxic drugs (P<4.9e-5, Supplementary Figure S2). Importantly, among the patients with high predicted migration levels, the patients treated with cytoskeletal drugs have better survival than those that were not treated with these drugs (KM ΔAUC= −0.47, log rank P<4.38e-4, Figure 4d, Methods). Thus, cytoskeletal drugs are more likely administered to patients with high (predicted) migration, and such administration seems to preferentially benefit such patients. This suggests that the predicted migration levels may serve as a biomarker for cytoskeletal drugs. As controls, patients with high predicted proliferation levels do not have a significant survival benefit from taking cytoskeletal drugs (KM ΔAUC= −0.2, log rank P<7.06e-2, Figure 4e). Finally, and interestingly, neither migration nor proliferation are good predictors of cytotoxic efficacy, as predicted migratory and proliferation levels are not associated with the survival of patients who have taken only cytotoxic drugs: (KM ΔAUC= −0.29, log rank P<1.08e-1, Figure 4f) for patients with high predicted migratory levels and (KM ΔAUC= −0.16, log rank P<1.38e-1, Figure 4g) for patients with high proliferation levels.

## Discussion

We experimentally measured migration and proliferation values in breast cancer cell lines and built and cross-validated CellToPhenotype predictors of these phenotypes. Applying these predictors to breast cancer tumor gene expression data we predicted the migration and proliferation levels of every tumor and studied their association with cancer stage, grade, subtype, and most importantly, with patient survival. We find a stronger association of predicted migration vs proliferation levels with the patients’ survival. We also find that patients with high predicted migration levels respond better to cytoskeletal drugs than patients with low predicted levels. siRNA-based predictors of migration and proliferation that we additionally built further testify that migration is indeed a better predictor of survival and the association may even have a causal basis. To the best of our knowledge, this is the first study which aims to quantify migration and proliferation levels in cancer patients by collecting and analyzing pertaining *in vitro* data. Such an investigation is particularly relevant since the majority of cancer drugs are developed by measuring their effect on *in vitro* proliferation rates^10^.

Many of the genes identified as migratory signatures by CellToPhenotype predictors are known to be play role in cell migration. For instance, the LAMA2 gene produces an extracellular protein, Laminin, is thought to play a role in migration and organization of cells in tissues during embryonic development^28^. CX3CR1 is known to play a role in adhesion and migration of leukocytes^29^. CX3CR1 is also amongst the top hits in our siRNA-based. RNT4 is another gene that is associated with cell migration^30, 31^. Our migration and proliferation signatures (either CellToPhenotype or siRNA-based analysis) identified many additional genes (Table S3p) that may be important in migration/proliferation, and their investigation may provide future leads for enhancing our understanding of these cellular phenotypes.

In summary, our analysis highlights the importance of tumor migration in determining its aggressiveness and patients’ survival. It puts forward the need to put more effort on *in vitro* assays of cell migration (and possibly, invasion) in the early stages of cancer drug development screens, which are yet mainly focused on essentiality and proliferation screens.

## Methods

### Quantification of cell migration and proliferation of a panel of breast cancer cell lines

#### Doubling times

Breast cancer cell line population doublings were estimated by plating a known number of cells at day 0 and measuring the total number of cells once the culture reached an estimated 80% confluency (usually 4-5 days). Cell numbers were calculated using a hemocytometer and population doublings (PDL) were determined using the following formula: PDL= 3.32 (log(total cells at harvest/total cells plated at day 0)). The doubling time was calculated by dividing the number of days or hours between harvest and seeding by the PDL. Example: Number of cells plated at day 0 = 5×10^6^, Number of cells at harvest = 20×10^6^, days in culture = 4 days, then the doubling time would be 2 days or 48 hours (4 days/2 PDL). The proliferation rate for each cell lines was computed using the equation 70/(doubling time). We did this for 46 breast cancer cell lines.

#### Live Cell migration assay^32^

Cells were seeded on 96-well glass bottom plates (Greiner Bio-one, Monroe, NC, USA) coated with 10 μg/ml collagen type I (isolated from rat tails) in PBS for 1 hour at 37°C. Before imaging, the cells were pre-exposed for 30-45 min to 0.1 μg/ml Hoechst 33342 (Fisher Scientific, Hampton, NH, USA) to visualize the nuclei. The plates were placed on a Nikon Eclipse TE2000-E microscope fitted with a 37°C incubation chamber and 5% CO_2_ supplier, a 20x objective (0.75 NA, 1.00 WD), an automated stage and perfect focus system. Up to four positions per well were automatically defined and nuclei (stained with live Hoechst) were imaged overnight every 10 to 20 minutes using NIS controlling software (Nikon) and a CCD camera (Pixel size: 0.78 or 0.32 μm). The .nd2 files acquired from NIS were exported to .tiff files as mono image for each channel and then converted to .avi files and analyzed using custom made ImagePro Plus macros as previously described^33^. The mean speed of cell migration was quantified per time-lapse by tracking each nucleus separately over time. This was done for 43 breast cancer cell lines.

### siRNA-image based migration assay using the MDA-MB-231 cell line^34^

#### Transient siRNA-mediated gene knockdown

Human siRNA of 475 protein kinases (whose gene expressions are significantly correlated with experimentally determined migration values in 40 breast cancer cell lines studied above, Table S2c) were purchased in siGENOME format from Dharmacon (Dharmacon, Lafayette, CO, USA). Transient siRNA knockdown was achieved by reverse transfection of 50 nM single or SMARTpool siRNA in 2,500-5,000 cells/well in a 96-well plate format (PKT assay) using the transfection reagent INTERFERin (Polyplus, Illkirch, France) according to the manufacturer’s guidelines. The medium was refreshed after 20 h and transfected cells were used for various assays between 65 to 72 h after transfection.

#### Phagokinetic track (PKT) assay

PKT assays were performed as described before^35^. Briefly, black 96-well μClear plates (Greiner Bio-One, Frickenhausen, Germany) were coated with 10 μg/ml fibronectin (Sigma-Aldrich, Zwijndrecht, The Netherlands) for 1 h at 37°C. Plates were washed twice with PBS, using a HydroFlex platewasher (Tecan, Männedorf, Switzerland). Subsequently, the plates were coated with white carboxylate modified latex beads (400 nm, 3.25 109 particles per well; Life Technologies, Carlsbad, CA, USA) for 1 h at 37°C, after which the plate was washed 7 times with PBS. 65 h after siRNA transfection, transfected cells were washed twice with PBS-EDTA and trypsinized. Cells were resuspended into single cell suspensions, then diluted, and finally seeded at low density (~100 cells/well) in the beads-coated plate. Cells were allowed to migrate for 7 h, after which the cells were fixed for 10 min with 4% formaldehyde and washed twice with PBS. For each transfection, duplicate bead plates were generated (technical replicates); transfection of each siRNA library was also performed in duplicate (independent biological replicate). Procedures for transfection, medium refreshment and PKT assay were optimized for laboratory automation by a liquid-handling robot (BioMek FX, Beckman Coulter).

#### PKT imaging and analysis

Migratory tracks were visualized by acquiring whole well montages (6×6 images) on a BD Pathway 855 BioImager (BD Biosciences, Franklin Lakes, NJ, USA) using transmitted light and a 10x objective (0.40 NA). A Twister II robotic microplate handler (Caliper Life Sciences, Hopkinton, MA, USA) was used for automated imaging of multiple plates. Montages were analyzed using WIS PhagoTracker20. Migratory tracks without cells or with more than 1 cell were excluded during image analysis. The quantitative output of PhagoTracker was further analyzed using KNIME. Wells with <10 accepted tracks were excluded. Next, data was normalized to mock to obtain a robust Z-score for each treatment and each parameter. After normalization, an average Z-score of the 4 replicates was calculated. Knockdowns with <3 images were removed, as well as knockdowns with <150 accepted tracks. Major Axis score (Z-score) as a measure of cell speed was further used and computed in the modeling.

### CellToPhenotype predictors

CellToPhenotype predictors consists of two expressions based supervised predictors – one for predicting cell proliferation and other for cell migration. The gene expression data was obtained from the Cancer Cell Line Encyclopedia project^36^. Each predictor was trained on in vitro cell migration or proliferation as the dependent variable and gene expression of cell lines as the independent variables in the regression. CellToPhenotype adopts two level feature selection to reduce testing error. First genes that are significantly associated with patient survival (in an independent dataset – METABRIC) were selected to be included in the subsequent regression model. Secondly, CellToPhenotype uses LASSO shrinkage to regularize the predictor that enables a data-driven feature selection using a cross-validation. Both above feature selection was conducted in dataset independent of the testing set on which performance of CellToPhenotye was evaluated. This ensures an unbiased evaluation of predictive power of CellToPhenotype.

To achieve the robust final estimate of migration and proliferation, CellToPhenotype uses bootstrapping. CellToPhenotype predictors were conducted on the training data and then phenotypes were predicted for test samples. The process is repeated for 50 bootstraps. Median of each bootstrap are taken as the final estimates of migration and proliferation levels (details are provided in the supplementary note).

### CellToPhenotype predictive performance using cross-validation

Leave one out cross validation was conducted to assess the CellToPhenotype predictive power as follows. The migration and proliferation models were trained on all in vitro data leaving one sample. Migration and proliferation was estimated for the left-out sample. Spearman ρ between the predicted and actual phenotypes was computed.

### Association of cellular phenotypes with cancer stages

Using CellToPhenotype predictors, migration and proliferation levels were predicted for 937 breast cancer patients in TCGA (Table S2a). Out of this, 83 and 9 individuals are stage IA and stage IB respectively (grouped as stage I); 348 and 238 individuals are stages IIA and IIB respectively (grouped as stage II); 148, 29, 62, and 19 individuals in stages IIIA, IIIB, IIIC, and IV respectively (grouped as stage III-IV). Patients with stage III and IV were grouped because there is only 19 sample from stage IV. These predicted levels were used to check how they vary with stages.

### Association of cellular phenotypes with cancer grade

Using CellToPhenotype predictors, migration and proliferation levels were predicted for 1706 breast cancer patients whose cancer grade information was available in the METABRIC dataset (146 grade 1 patients, 673 grade 2 patients, 887 grade 3 patients, Table S2a). These predicted levels were then used to check how they vary with grade.

### Association of cellular phenotypes with breast cancer subtypes

Migration and Proliferation levels are predicted for the 497 breast cancer patients (Table S2a) in TCGA dataset for whom we have the four different breast cancer subtypes information available, and of the 110 normal non-cancerous breast samples. A one-sided Wilcoxon rank-sum test was used to compare migration and proliferation levels of each of the subtypes with that of the normal samples.

### Circulating tumor cells analysis

We applied CellToPhenotype predictors to estimate the migration and proliferation levels in 5 breast cancer samples of circulating tumor cells (CTCs) in GSE45965 data^24^, and compared it with the 110 normal breast samples and 1043 cancerous samples from breast cancer TCGA data. While predicting migration and proliferation levels in GSE45965 data, we overlapped the survival associated genes in METABRIC dataset with the genes in the GSE45965 data, for building models using CellToPhenotype predictors.

### Association of cellular phenotypes with patient survival

Migration and proliferation models were built by training on 40 breast cancer cell lines that have experimentally measured migration and proliferation. Migration and proliferation levels of 1043 TCGA breast cancer patients were estimated using CellToPhenotype predictors. To check the association of the predicted migration with patients’ survival we fit following Cox regression:

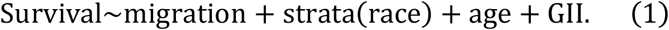

Patient survival is known to be confounded by age, race, and genomic instability (GII). Accordingly, the above model systematically controls for these confounders. Strata (race) in the above model implies Cox regression was conducted in each patient stratification based on race separately and likelihood were combined. We repeated the procedure for 10 iterations, and median coefficients (risk factor) of migration were computed. The association of survival with proliferation was estimated similarly. Each Kaplan Myer (KM) analysis was done by comparing the migration/proliferation levels of on the top 25 percentile of patients with bottom 25 percentile patients.

To estimate the relative contribution of migration and proliferation to predict patient survival we fit following Cox regression, which also controls for age, race, and genomic instability:

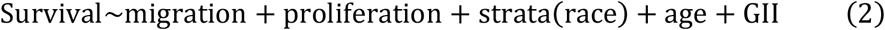

### siRNA-based KD-migration-score of a sample

Out of around ~4600 kinases, we selected 475 protein kinases whose expression was significantly correlated with migration across 9 cell lines (Spearman ρ, P < 0.01). Out of this, 248 protein kinases are negatively correlated and 227 protein kinases are positively correlated. We conducted siRNA knockdown of 475 genes described above in MDA-MB-231 breast cancer cell line. MDA-MB-231 was chosen because it is highly migratory breast cancer cell line^37^. Following the siRNA, we experimental measured change cell migration by measuring factors including Major Axis score (Z-score). Migration-suppressive genes (n=26) were identified by selecting genes whose knockdown led to high migration (above 90 percentile) in MDA-MB-231 cell line and whose gene expression was significantly negatively correlated with experimentally determined migration values in 40 cell lines. Migration-enhancer genes (n=24) are those whose knockdown significantly decreased cellular migration (below 10 percentile) and whose gene expression was significantly positively correlated with experimentally determined migration values in 40 cell lines. We count the number of downregulated migration-suppressive genes and upregulated migration-enhancer genes; and assign the count as KD-migration-score of each sample (cell line or patient). For this analysis, we use the median expression of all genes as the threshold for upregulation or downregulation.

### siRNA/shRNA-based KD-proliferation-score of a sample

siRNA/shRNA-based KD-proliferation-score of a sample is determined in an analogous manner described for migration. Briefly, we used proliferation measurement post ~15400 genes shRNA knockout in MDA-MB-231 cell lines from Marcotte *et al*.^25^. Among these genes, we select 1248 genes whose expressions are significantly correlated with experimentally determined proliferation values in 40 BC cell lines (606 genes positively correlated and 642 genes negatively correlated). Proliferation-suppressive genes were identified by selecting genes whose knockdown led to high proliferation (above 90 percentile) in MDA-MB-231 cell line and whose gene expression was significantly negatively correlated with experimentally determined proliferation values in 40 cell lines. Proliferation-enhancer genes are those whose knockdown significantly decreased cellular proliferation (below 10 percentile) and whose gene expression was significantly positively correlated with experimentally determined proliferation values in 40 cell lines. Count of down-regulated proliferation suppressive and upregulated proliferation-enhancer in a sample was assigned as KD-proliferation-score of the sample.

### Drug response analysis

Drug response information is available for 720 TCGA breast cancer patients in TCGA: 389 patients administering at least one of the 7 cytoskeletal drugs, and 331 patients administering at least one of the 31 drugs targeting only proliferation (i.e., cytotoxic drugs, Table S2b). Among the 1043 breast cancer patients who have high migration levels (top 25 percentile), we do a KM analysis among patients administering cytoskeletal drugs and the rest (Figure 4d). compares top 25 percentile patient with high migration levels with the rest of the 1043 patients. Similar KM analysis was conducted on patient administering cytotoxic drugs.

### Pathway and GSEA enrichment analysis

We also carried out GSEA analysis^38, 39^ on the genes selected by the LASSO regression in the CellToPhenotype predictors. The annotated gene sets from the Molecular Signature Database was used for this analysis^40^. GSEA analysis was done separately on these sets of genes. Enriched gene sets with FDR q-value < 0.05 is shown (Table S3). Similarly, we carried out GSEA analysis on *migration-enhancer/suppressive* and *proliferation-enhancer/suppressive* genes.

For each iteration of the LASSO regression in the CellToPhenotype predictors, we did KEGG pathway analysis. We also did KEGG pathway analysis on genes selected from siRNA-based analysis.

### Association with immune-infiltrating cell types

We predicted migration and proliferation levels in breast cancer TCGA data and compared them with the estimated tumor-infiltrating immune cell types^41^. For this, patients were divided into two groups (top 25 or bottom 25 percentile) based on their estimated TAM abundance levels or B cell abundance levels. Wilcoxon rank-sum test was done to see if migration/proliferation levels difference between the two groups.

## Notes

The authors declare no potential conflicts of interest.

